# Defining the cellular complexity of the zebrafish bipotential gonad

**DOI:** 10.1101/2023.01.18.524593

**Authors:** Michelle E. Kossack, Lucy Tian, Kealyn Bowie, Jessica S. Plavicki

## Abstract

Zebrafish are routinely used to model reproductive development, function, and disease, yet we still lack a clear understanding of the fundamental steps that occur during early bipotential gonad development, including when endothelial cells, pericytes, and macrophage cells arrive at the bipotential gonad to support gonad growth and differentiation. Here, we use a combination of transgenic reporters and single-cell sequencing analyses to define the arrival of different critical cell types to the larval zebrafish gonad. We determined that blood initially reaches the gonad via a vessel formed from the swim bladder artery, which we have termed the gonadal artery. We find that vascular and lymphatic development occurs concurrently in the bipotential zebrafish gonad and our data suggest that similar to what has been observed in developing zebrafish embryos, lymphatic endothelial cells in the gonad may be derived from vascular endothelial cells. We mined preexisting sequencing data sets to determine whether ovarian pericytes had unique gene expression signatures. We identified 215 genes that were uniquely expressed in ovarian pericytes that were not expressed in larval pericytes. Similar to what has been shown in the mouse ovary, our data suggest that *pdgfrb*+ pericytes may support the migration of endothelial tip cells during ovarian angiogenesis. Using a macrophage-driven photoconvertible protein, we found that macrophage established a nascent resident population as early as 12 dpf and can be observed removing cellular material during gonadal differentiation. This foundational information demonstrates that the early bipotential gonad contains complex cellular interactions, which likely shape the health and function of the mature, differentiated gonad.

**Summary Sentence:** Delineating the complex cellular interactions between vascular and lymphatic endothelial cells, pericytes, and macrophage in the bipotential gonad is essential for understanding the differentiation and functioning of the mature gonad.

**Graphical Abstract:** 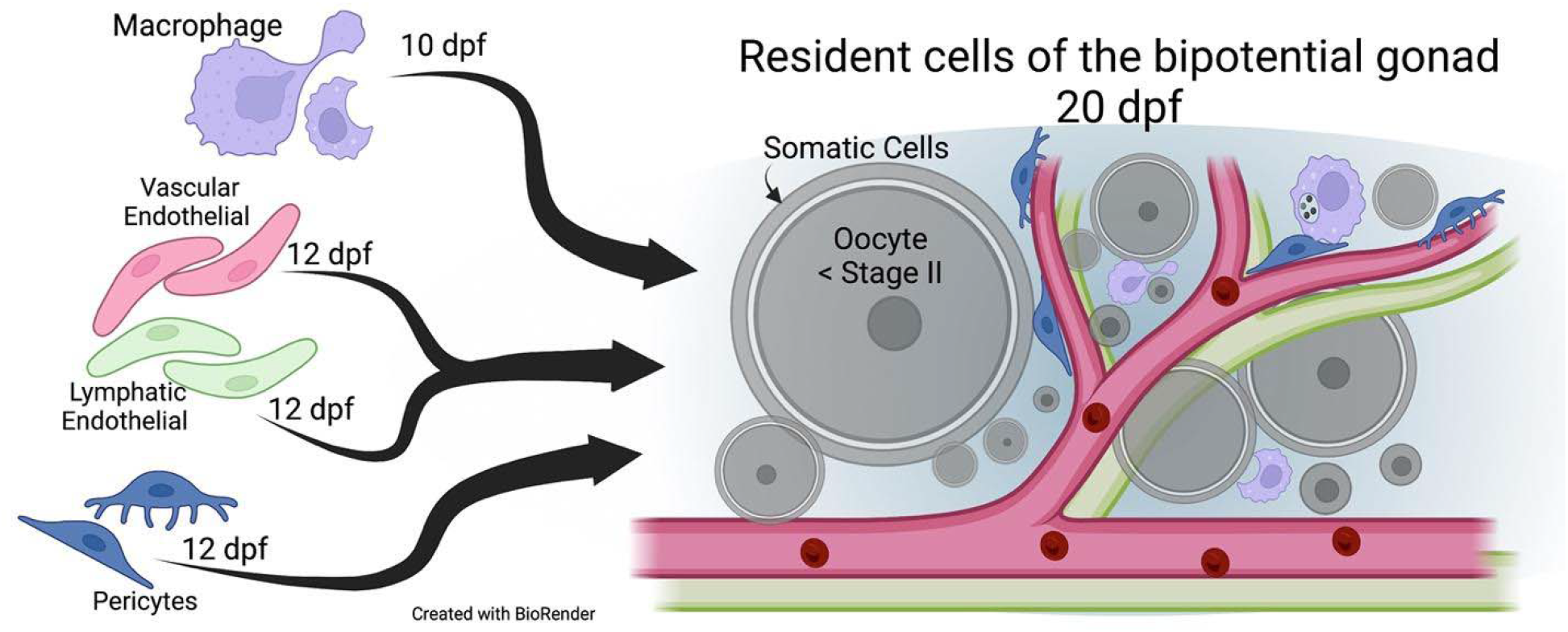

## Introduction

The developmental basis of the adult disease hypothesis postulates that the origins of some diseases can be traced back to developmental changes, sometimes very subtle, that produce long-term cascading effects that ultimately alter organ function and health (1). Given their genetic similarity to humans, zebrafish have been extensively used to model early embryonic development (2). More recently zebrafish have also been used to model juvenile development and adult health, including gonad formation and fertility (3–5).

The zebrafish gonad undergoes the same developmental stages as mammals; however, the timeline is elongated. The protracted developmental timeline allows for a detailed examination of very early periods of gonadal development, which have been understudied and may be sensitive to genetic and environmental disruptions. In mice and rats, primordial germ cells form at 7 days post conception (dpc) and, between ∼9.5 and ∼11.5 dpc, migrate from the hindgut to the urogenital ridge (6–8). In mice, the primary sex-determining gene, *sex-determining region Y* (*Sry*), must be activated before 11 dpc to initiate the transition from a bipotential gonad to a testicular development program (9). Since the murine bipotential period only lasts <2 days, it can be difficult to investigate early sources of reproductive disease that could occur during this critical developmental stage.

In mice, a major area of research has been to delineate the role of support and steroidogenic cells in sex determination and gonad maturation. Comparatively less has been done to investigate the role of other critical cell types in the developing bipotential and differentiated mammalian gonad. Research indicates that additional cell types, including macrophage, endothelial cells, and pericytes, are present at early stages of gonad development. At E10.5 in the mouse, macrophage are found at the urogenital ridge, just prior to sex determination (10). Macrophage depletion has been shown to hinder vascular remodeling and morphogenesis in the fetal testis (10). Endothelial cells in XX and XY individuals are present by 10.5 dpc and form lumenized vessels that are perfused with blood (10,11). Vessel formation is essential for testis development (12–14); however, considerably less is known about the dynamics of vascular development in the ovary as well as how lymphatic and perivascular cell development affects ovarian development (15). Recently, single-cell sequencing in 10.5 dpc mouse gonad and mesonephros found many different cell types are present at this early stage, including vascular and lymphatic endothelial cells, pericytes, and macrophage (16,17).

In the zebrafish, germ cells are specified at 4 hours post-fertilization (18). However, the primordial germ cells remain quiescent while migrating ventral of the hindgut, a process which occurs from 1 day post-fertilization (dpf) to approximately 10 dpf (see (19) for a detailed review). Somatic cells are shown in close association with primordial germ cells as early as 4 dpf (20). By 11 dpf, the somatic cells have formed a bilayer around the germ cells and begin to express sex-specific markers (21). From 10 dpf until approximately 20 dpf, most of the germ cells undergo mitosis, initiate meiosis, and remain in a bipotential state. During the bipotential period, while early-stage oocytes are forming, somatic cells surround the germ cells and the gonad exhibits mosaic expression of testicular- and ovarian-specific genes (i.e. anti-Mullerian hormone or aromatase, respectively). Early-stage oocytes in the bipotential period produce paracrine signals that act on somatic cells to further support oocyte development (21–23). The process of sex determination is initiated around 20 dpf. In the presence of sufficient oocyte-derived signaling, the oocytes continue to mature and, ultimately, form an ovary with mature granulosa and theca cells, which support ovarian health and function. In the absence of sufficient oocyte-derived signaling, oocytes undergo apoptosis and the gonad will progress with a testicular developmental program with Sertoli and Leydig cells supporting spermatogenesis (24,25). Thus, during the bipotential phase of zebrafish gonad development, although oocytes are present, the ultimate fate of the gonad is still undetermined. Unlike in the developing mouse testis and ovary, there is no mesonephric tissue adjacent to zebrafish gonad to give rise to the gonadal vasculature. There are still fundamental questions about the lineage of support cells in the differentiated ovary and testis. It is not definitively known if the somatic cells present during the bipotential phase are progenitors for mature support cells or if mature support cells of the adult gonad are derived from a different origin.

While it is known that the mature zebrafish ovary and testis contain endothelial and lymphatic vasculature as well as perivascular cells and macrophage (26), the developmental timeline for the arrival of these cell types to the gonad has not previously been described. The long bipotential period of gonadal development in zebrafish compared to mice provides an excellent opportunity to thoroughly assess the arrival of these critical cell types prior to sex determination. Using transgenic zebrafish with fluorescently expressed cellular markers, we tracked the arrival of the endothelial and lymphatic vasculature as well as perivascular cells and macrophage to the bipotential zebrafish gonad. We found that the bipotential gonad is more cellularly complex than originally thought. This foundational information suggests that additional cell types have the potential to contribute to gonad development, sex determination, and fertility.

## Material and Methods

### Zebrafish Rearing

Zebrafish (*Danio rerio*) were raised as described in Kossack et al. and Westerfield (27,28). Larvae were placed in tanks at 5 dpf in 100mL rotifer culture, 400mL of fish water static, and one drop of RG Complete (Reef Nutrition). From 5 to10 dpf, larvae were fed GEMMA Micro 75 (Skretting Zebrafish©) twice a day and live rotifers once a day. At 10 dpf, water flow was initiated as a slow drip with a rate of approximately one drop a second. From 10-20 dpf, fish were fed GEMMA Micro 75 twice a day along with rotifers and artemia once a day. From 21 to 28 dpf, the water flow rate was increased until a standard flow rate was established. The fish were treated as adults at 90 dpf and fed GEMMA Micro 300 once a day.

Juvenile fish were euthanized according to procedures approved by the Institutional Animal Care and Use Committee at Brown University and adhered to the National Institutes of Health “Guide for the Care and Use of Laboratory Animals”. In short, fish were placed in 0.04% MS-222 solution for 10-15 mins followed by ice water for 20 mins at which point they were decapitated and placed in 4% PFA overnight.

In establishing the developmental timeline, we measured the standard length of fish. The standard length is correlated with developmental stage during these ages (29,30). The full list of replicates, individuals, average length, and observations can be found in **Supplemental Table S1.**

### Transgenic lines utilized

To mark vascular endothelial cells, we used three transgenic lines, *Tg(fli1:nEGFP)^y7^, Tg(kdrl:GFP)* originally *Tg(flk1:GFP)^la116^,* and *Tg(kdrl:DsRed2)* (31–33). Lymphatic endothelial cells were labeled with *Tg(mrc1a:EGFP)* and *Tg(-5.2lyve1b:DsRed)* (*34,35*). *Tg(piwil1:EGFP)* originally *Tg(ziwi:EGFP)* marked germ cells, and *TgBAC(pdgfrb:EGFP)* marked perivascular cells (36,37). In the study of macrophage, we utilized *Tg(mpeg1:EGFP), Tg(mpeg1:Gal4FF)^gl25^,* and *Tg(UAS:Kaede)* (38,39). For readability and simplicity, we will refer to these transgenic lines by the simplified notations inside the parentheses, i.e *Tg(-5.2lyve1b:DsRed2*) is listed as *lyve1b:DsRed*.

### Antibody Staining

Antibody staining for germ cell protein Vasa was performed as described in Leerberg et al. (21) using anti-Vasa antibody from GeneTex (GTX1238306) at a working concentration of 1:1000. From 10-15 dpf, all gonads were left intact in the body cavity with the gut and excess anterior and posterior portions of the fish dissected away. In fish >15 dpf the gonad was dissected out of the fish before staining. All secondary antibodies were used at 1:500 and are as follows: goat anti-rabbit IgG 488 (Thermo Fisher Scientific Cat# A-11034, RRID:AB_2576217) and goat anti-rabbit IgG 633 (Thermo Fisher Scientific Cat# A-21071, RRID:AB_2535732). After protein labeling, tissues (either whole gonad or body cavity) were stained with Hoechst 33342 (Invitrogen H3570) at a concentration of 1:10,000 overnight, dissected, and placed in Vectashield mounting media (Fisher Scientific, NC9265087) for subsequent imaging.

### Photoconversion and live imaging

Double transgenic line *mpeg1:GAL4;UAS:Kaede* fish were identified at 3 dpf. At 12 dpf, fish were anesthetized in 0.02% MS222 for 1 minute then mounted on a 35mm glass bottom microwell dish (MatTek, Part No. P35G-1.5-14-C) in 2% low-melting temperature agarose (Fisher Scientific, bp1360-100) made in egg water (4 parts per thousand salt). The mounted fish was surrounded in 0.02% MS-222 and placed on the confocal microscope. Using 20x objective lens, the mid-section of the fish was exposed to full-power LED 405nm fluorescent light for 60 seconds to photoconvert the Kaede from green to red. The fish was immediately removed from agarose and replaced in the tank. The tank was covered with foil to protect the fish from additional light during transportation back to the fish facility. 24 hours later, the fish were again anesthetized and mounted laterally in agarose. Using a 20x objective lens, *piwil1:EGFP* expressing fish were identified from the previously photo-converted fish and imaged.

### Imaging

Images were collected on a Zeiss LSM 880 confocal microscope. Maximum intensity projections, orthogonal slices, and colocalization analysis were performed with ZenBlack (Zeiss). Tissues from 10 – 15 dpf zebrafish were mounted on Premium microscope slides (Fisher Scientific, 12-544-7) with Premium cover glass (Fisher Scientific,12-548-B) held in place by vacuum grease. Tissues >20 dpf were mounted in 1% low-melting agarose in a 35mm glass bottom microwell dish. All raw imaging files are available (https://doi.org/10.26300/4ce5-jk07).

### Sequencing Analysis

From Liu et al., (26) we obtained matrixes of gene and feature count. These large matrixes gathered from raw single-cell sequencing data display individual cells on one axis vs. counts of every gene detected on the other. From this data, we are able to perform principal component analysis to group similar cells together, which is referred to as clustering. For more details about the analysis see Seurat (40,41), and R code (https://doi.org/10.26300/4ce5-jk07). Clustering of the ovarian cells was performed as described in Liu et al. and cluster 5 was identified as the “Blood Vessel” cluster as described in the publication (26). We isolated this group using the subset() function and looked at the expression of vascular endothelial cell and lymphatic endothelial cell markers. We attempted sub-clustering analysis, but there were insufficient cell numbers within this cluster alone to draw meaningful information.

To generate the pericyte Venn diagram, we used publicly available data from Liu et al. and Shih et al. (26,42). Briefly, both publications included lists of highly expressed genes identified in all cell types. We, therefore, were able to download the list of genes expressed in the cell types of interest, filter them for expression level, and compare the lists. From Liu et al. (26), we downloaded the dataset of differentially expressed genes in “stromal cells” (from Supplementary file 5). Based on Lui et al. ovarian stromal cells are cells that are not able to be categorized as the following: germ, somatic, endothelial, or immune. Within ovarian stromal cells, there were 6 sub-types of cells: interstitial, pericytes, vascular smooth muscle, stromal progenitors, and ovarian cavity epithelial cells. From this data we created two groups, “ovarian pericytes” which included genes highly expressed in the pericyte sub-cluster 2, and “ovarian stromal cells” which included genes enriched in interstitial, vascular smooth muscle, stromal progenitors, and ovarian cavity epithelial cells (sub-clusters 0, 1, 3, 4, 5, respectively). We eliminated duplicates within each list to generate 1,831 genes in the ovarian stromal cell group and 441 genes in the ovarian pericyte group.

Similarly, using publicly available data from Shih et al. (42), we downloaded the highly expressed genes identified in all *pdgfrb*-expressing cells (from Table S2.1). As described in the publication we filtered for expression level log_2_>1, this list of genes was defined as “*pdgfrb*+ cells”. We performed the same methodology with Table S1.4, identified as “cluster 39”, which represents genes enriched in larval pericytes. We eliminated duplicates within each list, which resulted in 2,084 genes in *pdgfrb*+ cells and 105 genes in cluster 39. Using the Multiple List Comparator tool on molbiotools© (https://molbiotools.com/listcompare.php) we compared the groups and created a 4-way venn diagram. List details are available in **Supplemental Table S2**.

## Results

### Vascular endothelial cells make initial contact with the anterior gonad

To establish a timeline of vascular development in the gonad, we used a double transgenic line *kdrl:DsRed; fli1:nEGFP* that carries both the pan-endothelial marker *fli1:nEGFP* as well as *kdrl:DsRed*, which marks vascular endothelial cells. Beginning at 10 dpf, we used a qualitative scale to measure the distance between the gonad, marked by germ cells, and the vascular endothelial cells. We defined the relationship between the cells using three categories, “No Contact” (**Figure 1A**), “Proximal” (near, but not physically associated, **Figure 1B**, **Movie 1**), or “Contact” (direct physical contact, **Figure 1D, Supplemental Figure S1, Movie 2**). Because zebrafish do not have a coelomic epithelium, we defined the gonad as the cells within the boundary created by the somatic cells. We found that in most individuals (n=15/17), contact between vascular endothelial cells and germ cells occurred by approximately 20 dpf (**Figure 2A**), which corresponded to a standard length of ∼7mm (**Figure 2B, Supplemental Table S1**). We observed that vascular endothelial cells initially made contact with the anterior-most portion of the developing gonad (**Figure 1E, Movies 1 and 2**). Prior to this study, it was not known which vessels supplied blood to the bipotential gonad. We determined that at 20 dpf the blood supply to the bipotential gonad originates from the swim bladder artery (**Figure 1H**). Posterior to the anterior chamber of the swim bladder, the swim bladder artery divides, with two different branches supplying blood to the right and left gonad (**Figure 1H**). We termed this artery the gonadal artery.

**Figure 1.**
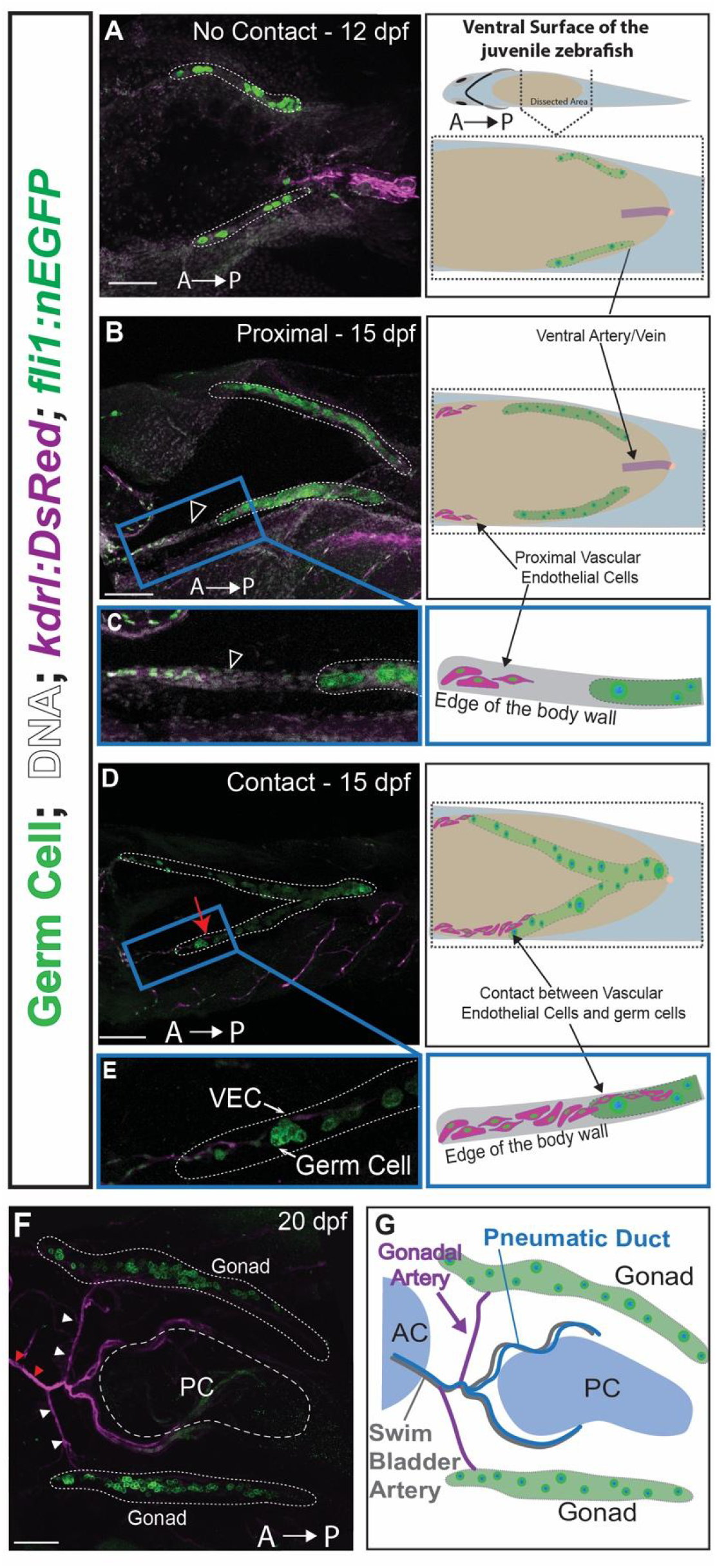
Vascular endothelial cells in the bipotential gonad. **(A-E)** Representative images showing the defined 3 categories for describing the relationship between vascular endothelial cells and germ cells in the biopotential gonad: “No Contact” (**A**), “Proximal” (**B-C**; open arrowhead, Movie 1), and “Contact” (**D-E**; red arrow, Movie 2). All images are oriented with anterior (A) to the left and posterior (P) to the right. (**F**) The main vessel supplying blood to the gonad, the gonadal artery (white arrowheads), originates from the swim bladder artery (red arrowheads). (**G**) A schematic representing the GA and SBA in the context of the body cavity. VEC = Vascular Endothelial Cell, AC = Anterior Chamber, GA = Gonadal Artery, SBA = Swim Bladder Artery, PD = Pneumatic Duct, AC = Anterior Chamber, PC = Posterior Chamber. Scale = 100 µm.

**Figure 2.**
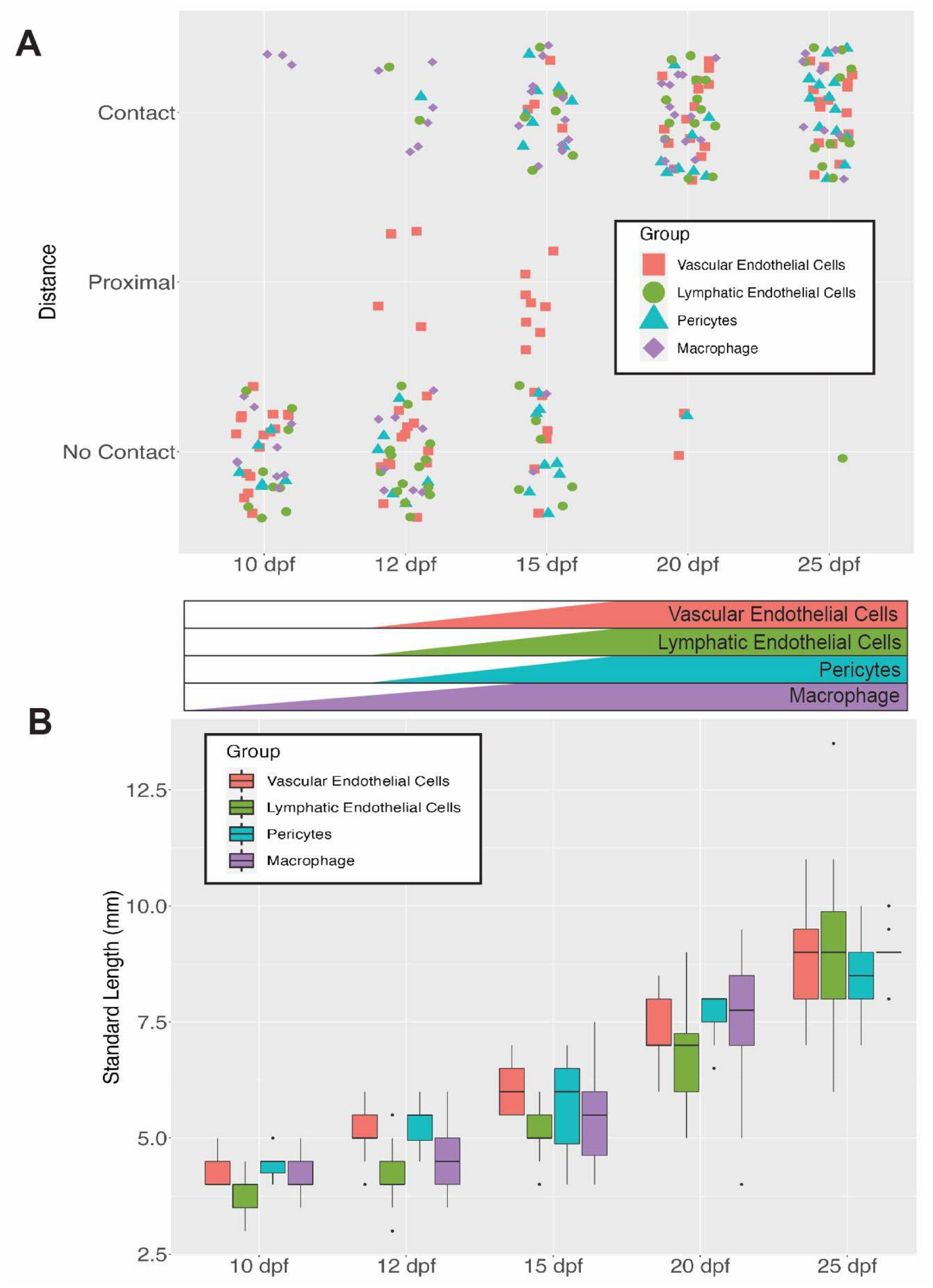
Vascular endothelial cells, lymphatic endothelial cells, and macrophage are present in the gonad when fish reach a standard length of 6.5mm. (A) At each age, individual fish from multiple spawns were collected to determine if there was contact between the cell of interest and the germ cells. Vascular endothelial cells (red circle), lymphatic endothelial cells (green triangle), pericytes (blue square), and sometimes macrophage (purple diamonds) arrive concurrently to the gonad between 15 and 20 dpf. (**B**) Each fish was measured prior to dissection to determine its length Separate cohorts of fish developed at different rates due to environmental conditions which led to variability in the initial point of contact. In juvenile fish, length can be used as a more accurate means of assessing developmental progression as compared to age. Contact between endothelial and germ cells occurred when the fish is approximately 6.5mm in length. The full list of replicates, individuals, average length, and observations can be found in **Supplemental Table S1**.

At 40 dpf, the ovary and testis are morphologically distinct. The ovary is significantly larger than the testis **(Figure 3A vs. 4A)**, however, both gonads are highly vascularized by this time point **(Figure 3B and 4B)**. By 60 dpf, each maturing oocyte appears to be surrounded by a vessel **(Figure 3E**, **Movie 3)**, while within the testis, vessels divide lobules of maturing sperm **(Figure 4E**, **Movie 4).** Within the ovary, we found that vessels have distinct relationships with different stage oocyte. Vessels come in direct contact with stage <1 oocyte, but are separated by a layer of somatic cells between the endothelium and the oocytes at stages 2 and 3 (**Supplemental Figure S2**).

**Figure 3.**
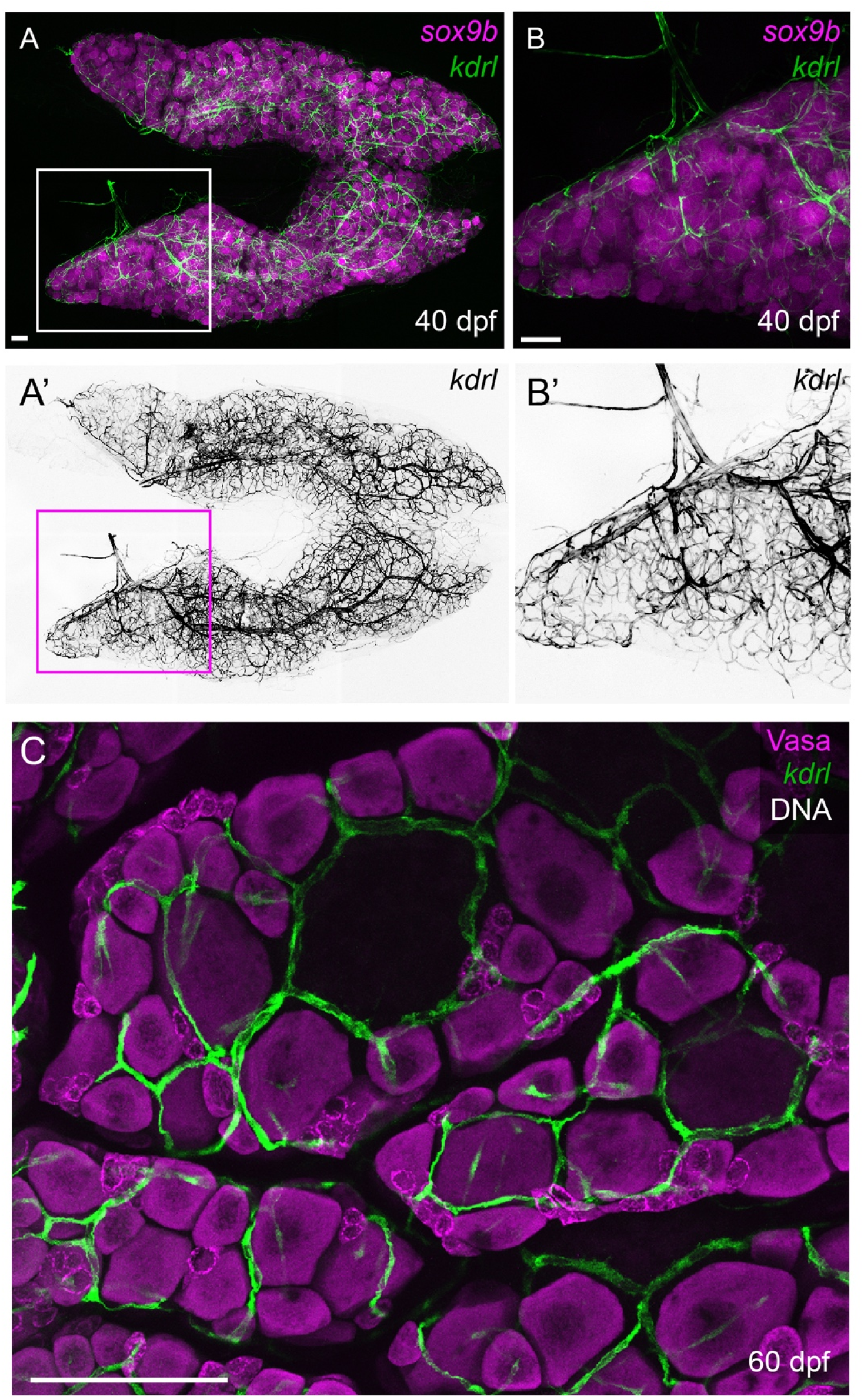
Vascularization in the zebrafish ovary. **(A-B’)** The two lobes of the zebrafish ovary are highly vascularized by 40 dpf. **(B-B’)** A high magnification of one ovary shows how large arteries quickly divide into smaller vessels surrounding oocytes. **(C)** As oocytes mature at 60 dpf, they will each be surrounded by at least one vessel (see Movie 3). *kdrl* and *sox9b* are, respectively, green and magenta in A, and B. *kdrl* is pseudo-colored black in A’ and B’. *kdrl*, Vasa, and DNA are, respectively, green, magenta, and grey in C. Scale = 100µm.

**Figure 4.**
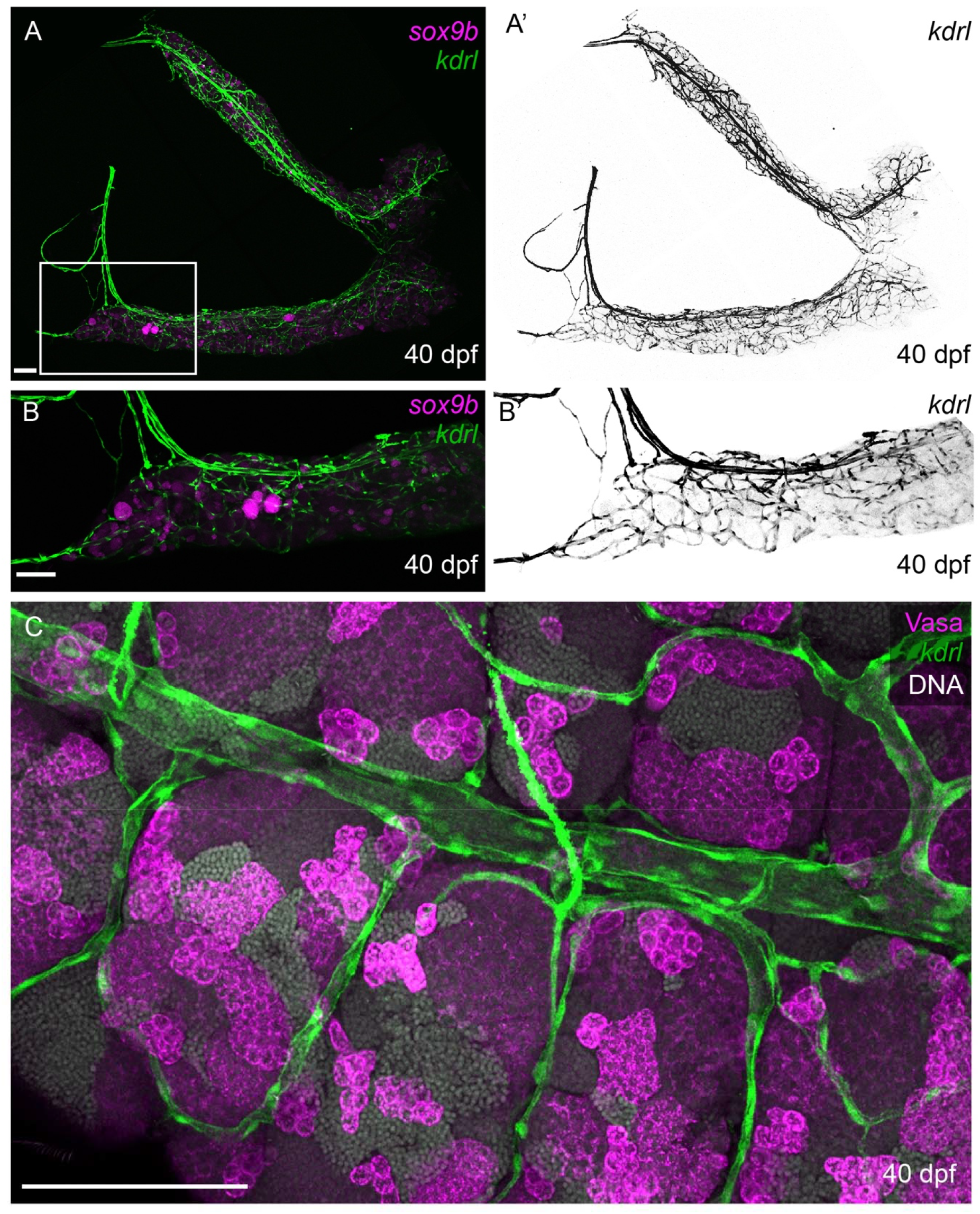
Vascularization in the zebrafish testis. **(A-B’)** The testis is significantly smaller than the ovary at 40 dpf, yet is also densely packed with vessels. **(B-B’)** High magnification of testicular vasculature at 40 dpf. **(C)** Vessels remain outside of the somatic cells of the 60 dpf testis and surround lobules of maturing sperm (see Movie 4). *kdrl* and *sox9b* are, respectively, green and magenta in A and B. *kdrl* is pseudo-colored black in A’ and B’. *kdrl*, Vasa, and DNA are, respectively, green, magenta, and grey in C. Scale = 100µm.

### Specification of lymphatic endothelial cells in the gonad resembles larval lymphatic development

The role of the lymphatic vasculature in organogenesis and homeostasis has been understudied in zebrafish (43). Recent single-cell sequencing of the zebrafish ovary found that lymphatic endothelial cells are present in the ovary by 40 dpf (26); however, it is not known when lymphatic cells invade the ovary and create functional networks within it. To determine when vascular and lymphatic endothelial cells reach the ovary, we visualized the endothelium by crossing canonical vascular endothelial reporter lines *kdrl:GFP* or *kdrl:DsRed* with established lymphatic endothelial transgenic reporter lines *mrc1a:EGFP* or *lyve1b:DsRed* (34). Using this double transgenic approach, we determined that lymphatic endothelial cells are detected in the gonad between 15 and 20 dpf and are always present with vascular endothelial cells (**Figure 2A**, 20 dpf, standard length of ∼7mm, n=13/13). Environmental conditions and genetic background will result in different growth rates; thus, the use of standard length is a more accurate measure of developmental stage and supports the concurrent arrival of VEC and LEC when the fish reach a length of ∼7mm (**Figure 2B**).

It is unclear from our initial analysis of early gonadal development if vascular endothelial cells give rise to lymphatic endothelial cells or if the lymphatic endothelium enters the gonad independent of the vascular endothelium. During larval development, lymphatic endothelial cells are derived from venous vascular endothelial cells (35,44–51). *flt4*, a marker of venous endothelial cells and lymphatic endothelial cells, is required for initiating lymphatic endothelial cell development. Expression of *flt4* is followed by the expression of *prox1a*, which specifies the lymphatic fate in mammals (51–58). However, in other contexts, such as anal fin vascularization, vascular endothelial cells are derived from lymphatic endothelial cells(57). To gain insight into which mechanism might be utilized in the gonad, we mined previously available single-cell sequencing from the 40 dpf zebrafish ovary (26). We sub-clustered the vasculature (“Blood Vessels”) and looked for expression and co-localization of specific endothelial markers. We found vascular endothelial markers *kdrl*, *flt1,* and pan-endothelial marker *fli1*, to be expressed in the majority of cells (**Figure 5A**) whereas *lyve1b*, *flt4*, and *prox1a* were expressed in a smaller subset of cells (**Figure 5B**). Next, we assessed whether *kdrl* and *lyve1b* were co-expressed in this cluster and we found no overlap in expression (**Figure 5C**). However, *lyve1b* was co-expressed with the venous marker *flt4*, suggesting that lymphatic development in the gonad may occur in a manner similar to what has been observed in the larval endothelium (34). We note that we were not able to examine the expression of *mrc1a* in the single-cell sequencing data set because it was not detected in the original sequencing analysis.

**Figure 5.**
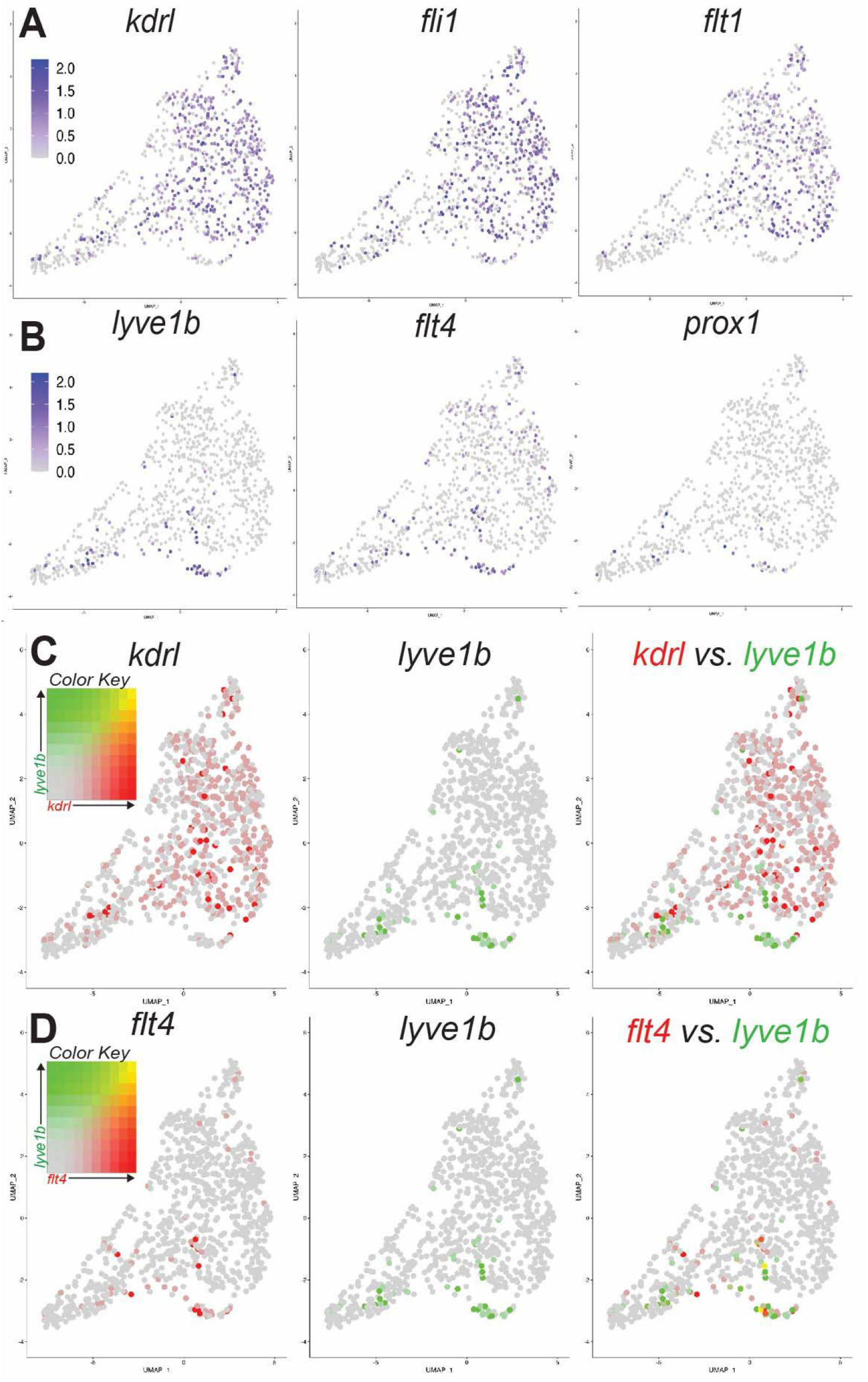
Endothelial gene expression in 40 dpf ovary. **(A)** Expression of vascular endothelial cell genes *kdrl, fli1*, and *flt1* occurred in a majority of the endothelial cells. (**B**) lymphatic endothelial genes *lyve1b*, *flt4*, and *prox1* were expressed in a smaller subset of cells. (**C**) There was no overlap in expression of *kdrl* (red) and *lyve1b* (green) by single-cell sequencing. (**D**) There was overlap in expression of *flt4* (red) and *lyve1b* (green) suggesting that lymphatic endothelial cells originate canonically.

To assess if lymphatic endothelial cells are derived from vascular endothelial cells in the bipotential gonad, we used confocal microscopy to image 20 dpf gonads expressing transgenic markers for both vascular endothelial cells and lymphatic endothelial cells (*kdrl:DsRed; mrc1a:EGFP*). Our imaging revealed overlap in the *kdrl:DsRed* and *mrc1a:EGFP* expression domains (**Figure 6A**). However, in most of the cells that expressed both genes, the level of transgenic expression was not equal. We consistently observed expression of one marker being dominant over the other in the maximum intensity projection (**Figure 6B and 6C**). Orthogonal slices of the z-stacked images illustrate that the incongruent levels of expression between markers are not an artifact of the maximum intensity projection (**Figure 6D**). Finally, we used Zen Black to perform a co-localization analysis of the confocal images and found that although the *mrc1a:*EGFP expression obscured the visibility of the *kdrl:DsRed*, there was 100% overlap in signal (**Figure 6E**, Overlap coefficient = 0.97). Together this suggests that lymphatic endothelial cells in the 20 dpf zebrafish gonad are derived from vascular endothelial cells, and as lymphatic specific expression increases, vascular endothelial expression decreases.

**Figure 6.**
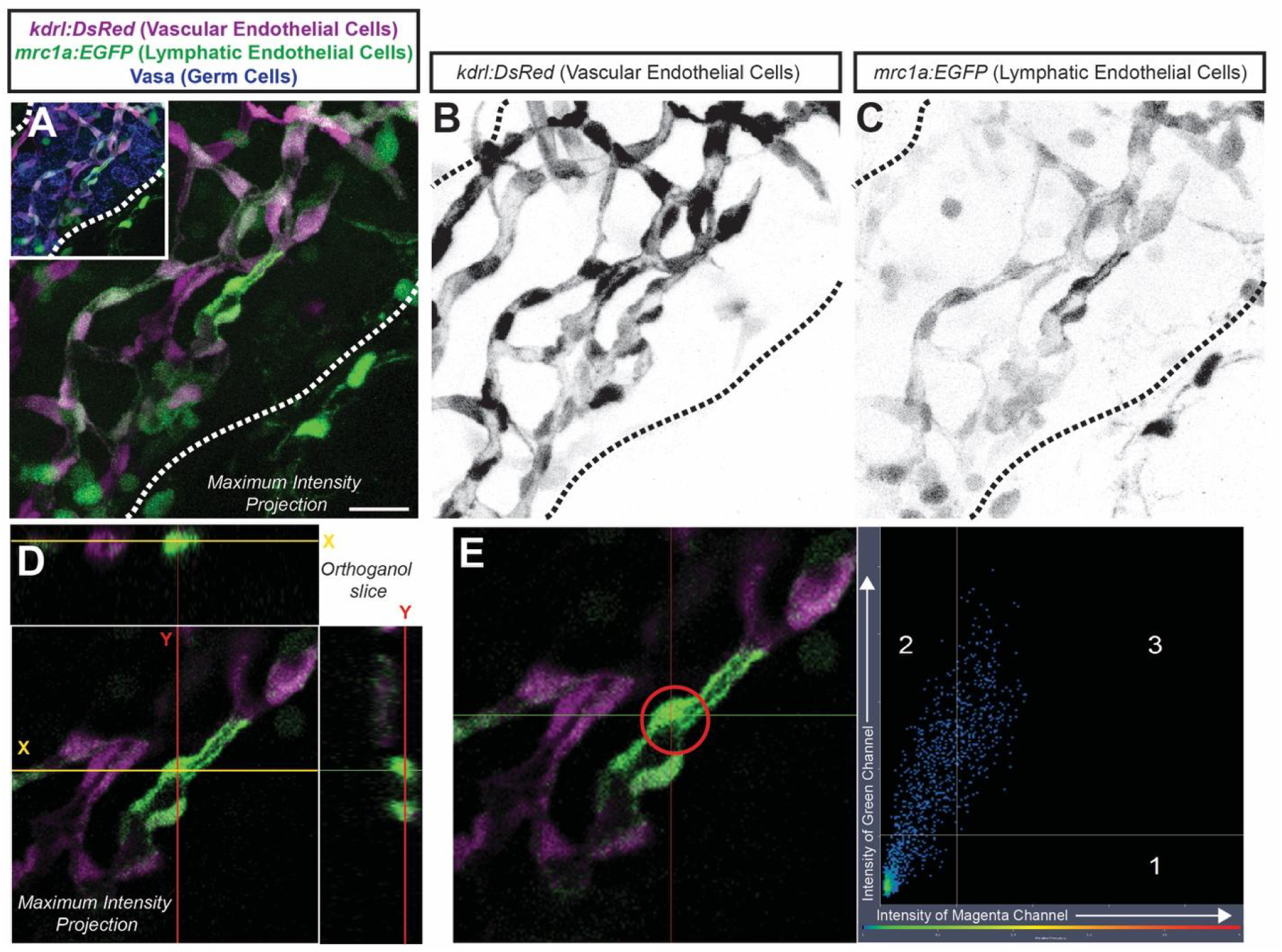
*kdrl* and *mrc1a* are co-expressed in the 20 dpf ovary. (**A**) Maximum intensity projection of the 20 dpf gonad (germ cells = blue, gonad outlined) showing overlap in *kdrl* (magenta) and *mrc1a* (green) expression. (**B**) Single channel showing *kdrl* expression alone suggests that *kdrl* is expressed in all cells. (**C**) Single channel showing *mrc1a* expression is inversely correlated with the strength of *kdrl* expression. (**D**) Orthogonal slice of the maximum intensity projection shows that the expression of both markers is overlapping within cells. (**E**) Co-localization analysis of the cells circled in red in (**F**) showing expression of *kdrl* and *mrc1a* is highly correlated. Scale = 20 µm.

### Pdgfrb+ perivascular cells associate with vascular tip cells invading the bipotential gonad

Pericytes are a sub-type of perivascular cells that have, since their discovery, been difficult to clearly distinguish from other perivascular cell types (59). In the developing zebrafish embryo, pericytes were originally defined by their expression of platelet derived growth factor receptor ß (*pdgfrb)*, their morphology, and their positional relationship to the vasculature. Recent single-cell sequencing data from larval zebrafish as well as the imaging of *pdgfrb*:EGFP zebrafish indicate that while pericytes tend to have high *pdgfrb* expression, they constitute only a subset of the *pdgfrb*:EGFP+ cells (42). The subset of *pdgfrb*:EGFP+ cells with high *pdgfrb* expression and low expression of other cell type markers was termed cluster 39 in a study by Shih et al., using single-cell RNA sequencing to examine the signatures of gene expression in larval zebrafish pericytes (42). Single-cell sequencing of the zebrafish ovary showed that *pdgfrb*+ cells are found in a cluster of “stromal cells” defined as non-germ, non-somatic, non-endothelial, and non-immune cells. Within the ovarian stromal cell cluster, ovarian pericytes had the highest *pdgfrb* and *notch3* expression, known pericyte markers in larval zebrafish, and were more uniquely defined by the expression of *plp1b* (26). We hypothesized that ovarian pericytes, as defined by Liu et al., and the larval pericyte cluster (cluster 39) would have more similar expression profiles than other *pdgfrb+* cells in the larvae or ovary. We compared expression profiles of larval *pdgfrb*:EGFP cells and the larval pericytes (cluster 39) from Shih et al., with ovarian stromal cells and ovarian pericytes from Liu et al. (26, 42). 215 genes were identified as unique to ovarian pericytes (a complete list of overlaps in **Supplemental Table S2**). This finding suggests that the 215 genes detected, including *plp1b*, may be unique to ovarian pericyte function and not required at larval stages of pericyte development. Only 3 genes, *pcdh1b*, *ptenb*, and *s1pr1* were found uniquely in ovarian pericytes and cluster 39 (**Figure 7**) *pcdh1b* and *s1pr1* are associated with cell adhesion (60,61) and are also both expressed in the follicle cell, germline stem cell, and endothelial cell clusters. *ptenb* is expressed broadly in all of the ovarian cell clusters and is a tumor suppressor gene known to control cell proliferation during embryogenesis however its functions in the zebrafish ovary is unclear (62). *notch3,* along with 10 other genes, was expressed in all four cell groups.

**Figure 7.**
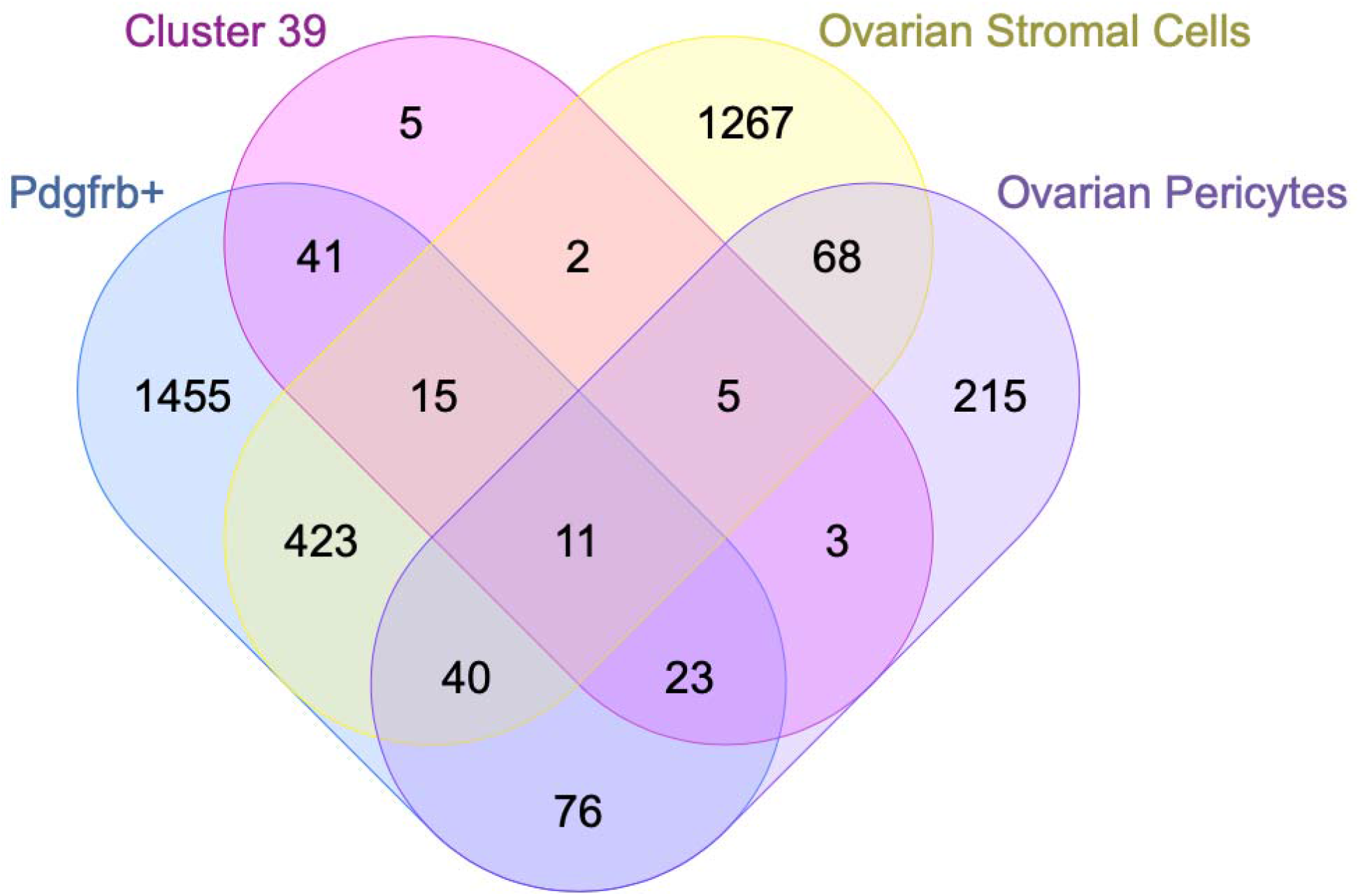
The *pdgfrb+* expression domain encompasses multiple cell types in the ovary. Using previously published data from single-cell sequencing in the embryo, we compared expression profiles of *pdgfrb*+ cells (blue) to cluster-39 (pink), which is a subset of *pdgfrb*+ cells with the highest *pdgfrb* expression and low expression of other known cell type markers. We further compared larval pericytes to 40 dpf ovarian stromal cells (yellow), which are defined as non-germ, non-somatic, non-immune, and non-endothelial cells, and ovary-specific pericytes (purple) using single-cell data from the zebrafish ovary. 215 genes, including *plp1b,* were found to be unique to ovarian pericytes (purple only), and 3 genes were expressed only in cluster 39 as well as ovary-specific pericytes (pink and purple overlap). *notch3* was expressed in all four cell groups (yellow, purple, pink, and blue overlap).

As with all vascular beds, pericytes in the ovary are an essential part of the vascular unit, creating stability for the constant vascular modeling that occurs as an inherent part of ovarian function (as reviewed in (63)). Since *pdgfrb* is an imperfect marker of pericytes, we used the double transgenic *pdgfrb:EGFP;kdrl:DsRed* to determine if *pdgfrb*+ cells within the gonad were also in contact with vessels. We found that oblong *pdgfrb+* cells are associated with vessels in the developing gonad and based on their gene expression, position, and morphology, appeared to be pericytes. The *pdgfrb+* presumptive pericytes arrived at the gonad along with the endothelial cells (**Figure 2A**, 20 dpf, n=8/9). Confocal images revealed *pdgfrb+* cells in close association with *kdrl+* vessels and extending tip cells (**Figure 8**, **Movie 5, Supplemental Figure S3**), suggesting that pericytes may aid in the migration of vascular tip cells, which has been observed during the vascularization of other tissues (37,64).

**Figure 8.**
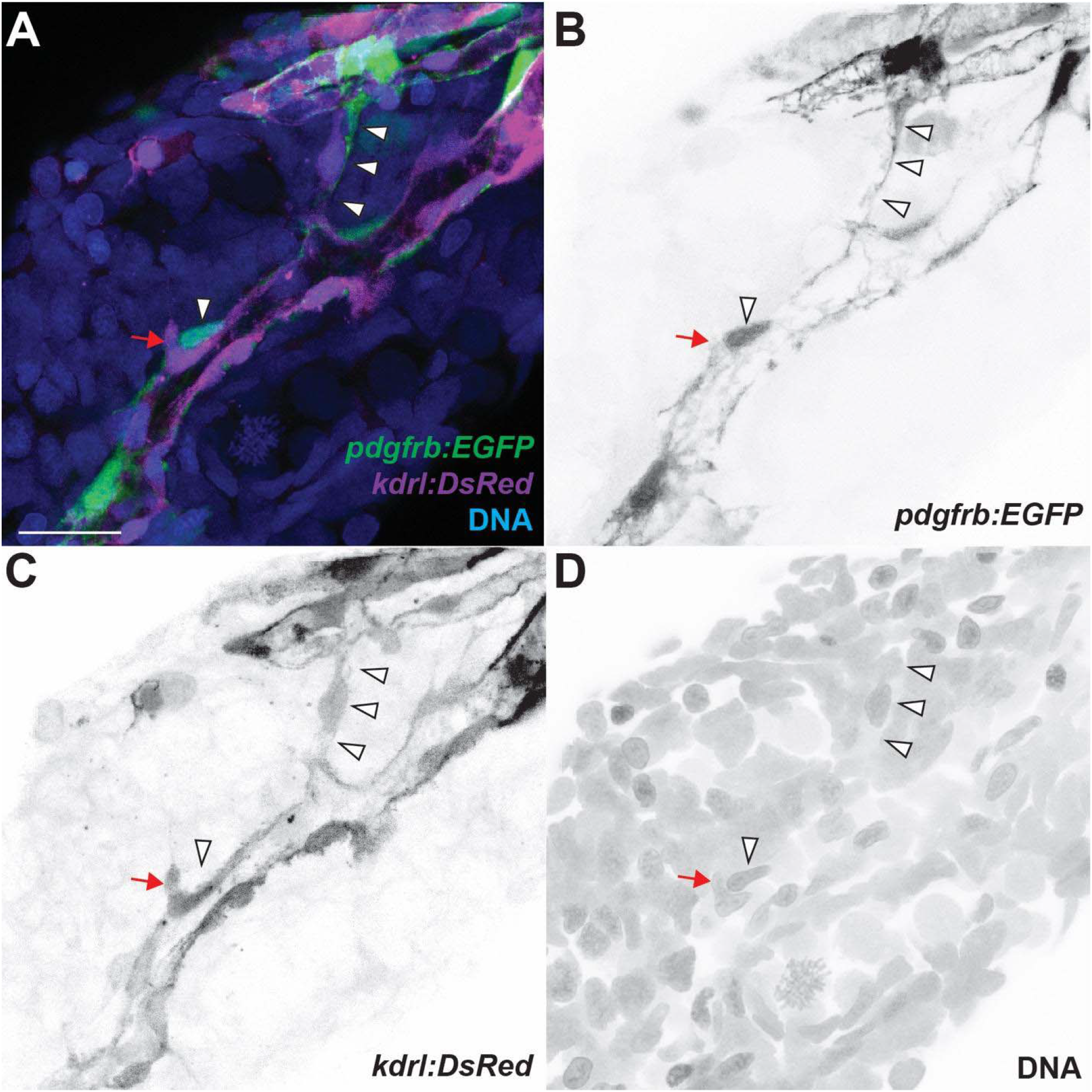
*pdgfrb* positive cells elongate in close association with vascular endothelial tip cells. (**A)** Maximum intensity projection of all channels of the 20 dpf gonad show pericytes, labeled with *pdgfrb:EGFP* (green). (**B**) Pericytes (pseudo-colored black) extend protrusions (arrowheads) between two existing vessels. *pdgfrb:EGFP* cells have classic pericyte morphology and are located at the division of vessels supporting the conclusion that they are pericytes (arrowheads). (**C**) *kdrl:DsRed* vessels (pseudo-colored black) are associated with an endothelial tip cell. (**D**) DNA (pseudo-colored black). See Movie 5. Scale = 20 µm.

### Macrophage arrive at the bipotential gonad prior to stromal cells types

Tissue resident macrophage play essential roles in organogenesis and organ homeostasis and macrophage are known to play a substantial role in both ovary and testis function in mice. However, the timeline of macrophage colonization of the zebrafish gonad remains unclear (65–67). We found that while macrophage colonization of the gonad in most individuals was concurrent with vascularization of the gonad, in some cases we observed macrophage within the gonad prior to vascularization (**Figure 2A**, 12 dpf n=6/14, 3 replicates). Notably, we found that at 20 dpf, when the gonad is undergoing sex determination, there are more macrophage present, and they can be seen engulfing germ cells (**Figure 9B, 9C, Movie 6**). This suggests that macrophage may play an active role in gonad differentiation in zebrafish.

**Figure 9.**
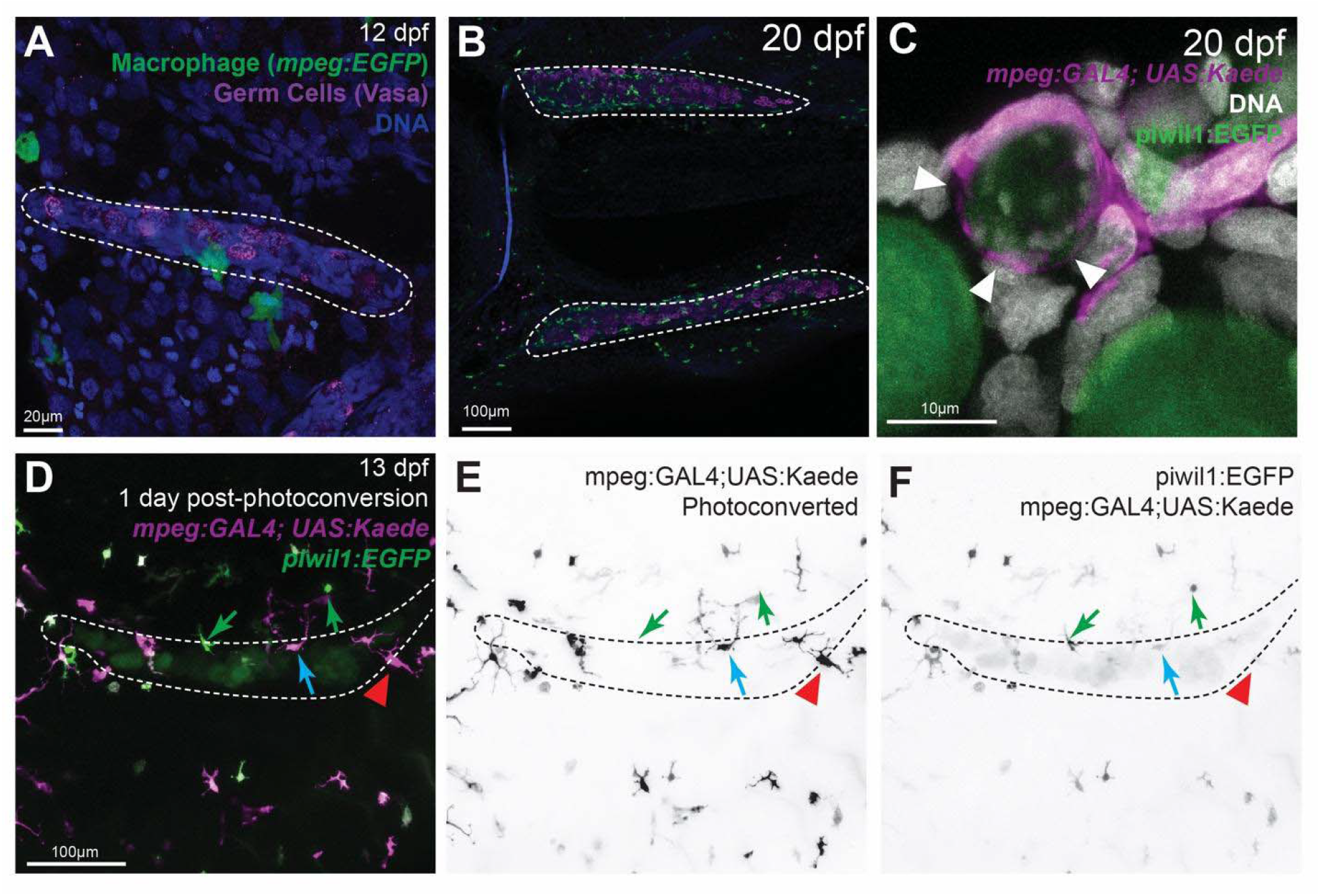
Tissue-resident macrophage engulf cells during gonad differentiation. (**A)** Macrophage were found in contact with germ cells as early as 12 dpf. (**B**) At 20 dpf, macrophage are abundant in the gonad. (**C**) Macrophage (magenta) were observed removing oocyte material (green) and DNA (white) from the former bipotential gonad (white arrowheads, Movie 6). (**D**) *mpeg1:GAL4; UAS:Kaede* expressing macrophage were photoconverted from green to red (pseudocolored magenta) at 12 dpf. 24 hours after conversion green and red macrophage could be found in the gonad (blue arrow, outlined). (**E**) Some macrophage in the gonad were only magenta (red arrowhead), indicating that they are resident to the gonad during this period. (**F**) In the green channel, we can observe that some macrophage migrate to the gonad within the 24 hour window post-photoconversion (green arrows). See Movie 7 and 8. Scale listed.

We asked if the macrophage that make initial contact with the gonad became resident macrophage or if they were only transiently interacting with the germ cells. Given the superficial position of the macrophage in contact with the early gonad (**Figure 9A**), we hypothesized that macrophage found during the bipotential stage were not tissue-resident and the resident population would not be established until after sex determination. To test this hypothesis, we used a pan-macrophage reporter to drive the expression of Kaede *(mpeg1:GAL4; UAS:Kaede)*, a photoconvertible fluorescent protein that changes from green to red following exposure to 405nm light. We exposed fish at 12 dpf to 405nm light in a defined region surrounding the gonad (**Supplemental Figure S4**). As a result, macrophage in and near the gonad were converted to red, leaving the remaining macrophage in the body green. 24 hours later we used live imaging to observe the macrophage population within the gonad. The presence of only red macrophage would indicate that no new macrophage had entered the gonad and suggest that macrophage become resident prior to sexual differentiation. The presence of green macrophage in addition to red macrophage would indicate either: (1) continued colonization of the gonad, (2) the presence of transient populations of macrophage, or (3) newly divided macrophage from a previously photoconverted cell although likely some red would remain from the mother cell. We found that after 24 hours, a subset of macrophage were retained in the gonad and expressed the red photoconverted kaede protein (**Movie 7 and 8**, **Figure 9E**, red arrowhead, n=5). We also observed green macrophage in and in the proximity of the gonad (**Figure 9F**, green arrow). Finally, several cells also were green and red mixed suggesting that they had some previous conversion while generating new Kaede protein (**Figure 9D**, blue arrow). It is difficult to say definitively where the double-labeled cells originated. Given the ratio of green to red in these cells, it is possible that these cells were on the periphery of the conversion area and migrated to the gonad while producing new Kaede protein. Together, these data suggest that at 12 dpf, there is an early population of resident macrophage that cells within the bipotential gonad.

## Discussion

Herein, we provide the first report of vascularization of the early zebrafish gonad, a fundamental and understudied component of reproductive development and health. We found that vascular endothelial cells and lymphatic endothelial cells are present in the ovary by 15 dpf. In mice, previous studies have tracked the arrival of endothelial cells to the early gonad using fluorescent imaging and single-cell sequencing at E10.5 (11,16,17). Based on single-cell sequencing from Liu et al (26), we know that a diverse population of cells are present in the 40 dpf zebrafish ovary. Yet without a developmental timeline, we lack key spatial and temporal information about gonadal development. Based on these previous studies, we can conclude that there are parallels between the vascularization of the zebrafish and mouse bipotential phase of the gonad. Although our investigation clearly indicates vascular and lymphatic endothelial cells are present in the bipotential gonad, it does not evaluate when the vessels establish a functional network. Future investigations will be required to determine when the vessels are lumenized and are able to transport physiological signals and waste.

Prior mouse studies did not rigorously study both vascular and lymphatic endothelial cell markers simultaneously in early development and, therefore, these studies do not allow us to distinguish between endothelial cell types during gonad development (15,68,69). In adult mice, vascular vs. lymphatic vessels have distinct domains in both the ovary and testis (11,69), but how these domains are established in early development is unclear. In mice, Brennen et al. (11) found that lymphatic endothelial cells, marked by Prox1^+/lacZ^, are present in the mesonephros at 13.5 dpc and enter the gonad at 17.5 dpc in both sexes. However, using a transgenic Prox1-EGFP mouse, Svingen et al. found that the ovary does not contain lymphatic endothelial cells until after birth at post-natal day 10, while the testis was reported to contain lymphatic endothelial cells by 17.5 dpc (69). Yet murine expression analysis using single-cell sequencing shows evidence of *prox1* expression in the gonad at 10.5 dpc (16,17). Each of these studies used different methods and criteria (lacZ, transgenic line, mRNA expression, respectively) to determine the presence of lymphatic endothelial cells in the gonad using *Prox1*. Therefore, the observed discrepancies could be due to differences in methodology or criteria. When comparing the zebrafish and mice bipotential gonad, it becomes more difficult to assess the parallels between lymphatic development in these two model species. Most murine studies use *Prox1* as the lymphatic endothelial cell marker. In our study, we defined lymphatic endothelial cells to be *mrc1a* or *lyve1b* expressing cells, because *prox1* is known to be expressed in a number of different ovarian cell types and is not required for lymphatic cell specification in zebrafish (49,54). Single-cell sequencing is able to detect extremely low expression of *Lyve1*, the mammalian orthologs of *lyve1b*, in endothelial cells at 10.5 dpc, which suggests the earlier timeline of lymphatic endothelial cell arrival to the gonad observed in zebrafish may also occur in mice (17). It is well established that lymphatic vessels are not found within the testis, but instead restricted to the outer tunicate; therefore, more research is needed to determine the organization of lymphatics in relation to the adult zebrafish testis and the differences between zebrafish and mouse sex-specific lymphatic development (10,69,70).

We also demonstrated that the *pdgfrb* expression domain within the ovary is not limited to pericytes and likely includes other perivascular cells with different sub-functionalizations. We show that *pdgfrb*+ cells extend between vessels within the gonad and are in close association with migrating vascular tip cells, a phenomenon that has been observed in other developmental contexts where vascular tip cells release *pdgf* ligands, which signal perivascular cells to follow and stabilize the forming vessel (37,64,71).

Pericyte biology is still relatively nascent and *pdgfrb* has been routinely used as the primary marker across species. In embryonic zebrafish, *pdgfrb* is required for the formation of the intersegmental vessels and meningeal angiogenesis (72,73). In other species, it is known that pericytes are essential for vascular stability and reconstruction of the ovarian vasculature during follicle development and post-ovulation (74–77). Consistent with this finding, inhibition of Pdgfrb in the rat ovary causes hemorrhaging (78). Together, these reports indicate a role of pericytes in the function and establishment of the blood follicle barrier (79,80). When we compared the gene expression profiles of *pdgrfb*+ cells with larval pericytes (cluster 39), ovarian stromal cells, and ovarian pericytes, we found that 215 genes were unique to ovarian pericytes, and only three genes were shared solely between ovarian pericytes and cluster 39. Of these, we note that two were associated with cell adhesion (*pcdh1b* and *s1pr1*). *s1pr1* has been shown to play critical roles in negatively regulating angiogenesis in zebrafish (60) and knockout of S1pr1 in mice causes size-selective openings in the blood-brain barrier (61). Although it is *s1pr1* is not specific to pericytes, as it is also expressed in other ovarian cells as well as in vascular endothelial cells including hepatocellular carcinoma (81, 82), the shared expression of cell adhesion genes in larval pericyte cluster (cluster 39) and ovarian pericytes supports the idea that pericytes play a critical role in the blood follicle barrier development in zebrafish. It will be necessary to understand how pericytes interact with the ovarian vasculature, particularly in addressing ovarian disease. Both ovarian hyperstimulation syndrome and polycystic ovary syndrome are associated with altered ovarian vasculature (83–85). Although it is well established that pericytes perform a critical role in vascularization and angiogenesis, the role that perivascular cells play in disease etiology remains unclear (reviewed in (84)). Zebrafish are able to model some, but not all aspects of ovarian diseases such as PCOS (86), therefore the discovery of additional cell types in the zebrafish bipotential gonad offers an opportunity to investigate the role of these cells in the very early reproductive disease formation.

We discovered that macrophage are present in the early bipotential gonad and can be seen engulfing germ cells during the bipotential phase. In mice, yolk-sac-derived macrophage are responsible for clearing cellular debris from endothelial cells, Sertoli cells, and germ cells, as well as for pruning vascular networks in the testis (10). In other contexts, macrophage are known to release angiogenic cues that shape vascular development (87–91). Therefore, their presence in the bipotential gonad prior to the arrival to vascular endothelial cells could suggest that macrophage are be involved in directing vascularization of the gonad. While ovarian resident macrophage are essential for folliculogenesis, ovulation, and the clearing of atretic follicles (67,92,93) as well as cellular debris (66), prior to this study, macrophage phagocytosis in the zebrafish ovary had not been observed. We also report that resident macrophage in the gonad are present by 12 dpf and the population is dynamic during this developmental window with new macrophage migrating into and out of the gonad. In mice, ovarian resident macrophages can be divided into 5 different populations. Two of these initial subpopulations are embryonically derived, one population from the yolk-sac, and the other from the fetal liver (66). However, the majority of research studies have investigated resident macrophage function after sex determination. The presence of macrophage in the zebrafish bipotential gonad suggests that macrophage may also be present in the mammalian bipotential gonad and contribute to important steps in sexual differentiation.

In conclusion, we found that vascular and lymphatic endothelial cells, pericytes, and macrophage begin populating the bipotential gonad at 12 dpf. Our data suggest that vascular endothelial cells may give rise to lymphatic endothelial cells in the gonad to together create a complex network of endothelial cells by 20 dpf. However, it is still unclear when these vessels become functional conduits for physiological signals and nutrients and additional loss of function experiments are necessary to confirm this hypothesis. We found that *pdgfrb+* pericytes support the migration of vascular tip cells and 215 genes are unique to ovarian pericytes compared to larval pericytes. Finally, we found that tissue-resident macrophage can be found in the gonad as early as 12 dpf and clear oocyte debris during sexual differentiation. Our foundational information regarding vascular and lymphatic endothelial cell, perivascular, and macrophage invasion into the biopotential gonad contributes to the understanding of the early bipotential environment and how they may play a part in the etiology of reproductive disease.

## Acknowledgments

We would like to thank Rachel Cyr for all her meticulous work caring and raising the fish. We would also like to that the Weinstein and Draper labs for sharing their transgenic lines that made this study possible. This work was made possible by funding from NIEHS to Dr. Michelle Kossack F32ES023650 and 5T32ES007272. Dr. Jessica Plavicki was supported by an NIEHS K99/R00 (ES023848), a CPVB Phase II COBRE (2PG20GM103652), and an NIEHS ONES award (ES030109).

## Data Availability

The data underlying this article are available in the Brown Digital Repository at: https://doi.org/10.26300/4ce5-jk07

## Author Contribution

MK: Conceptualization, methodology, investigation, writing – original draft, review, and editing, visualization. LT and KB: Investigation, writing – review and editing. JSP: Resources, writing – review and editing, supervision, project administration, fund acquisition.

## Movie Legends

**Movie 1: 3D rendering of “Proximal” association between vascular endothelial cells and germ cells.** In the center of the plane, the vascular endothelial cells elongate towards the germ cells at 12 dpf. DNA from either the somatic gonad or the body wall is present between the vessel and germ cells. *kdrl* (vascular endothelial cells, green), Vasa (germ cell, magenta), DNA (Blue).

**Movie 2: 3D rendering of “Contact” between vascular endothelial cells and germ cells from Figure 1D**. At ∼9 seconds the germ cells (green) contact with the vascular endothelial cells (magenta) are in close view. At ∼16 seconds, vessels running vertically are in view, these exist in the body wall between somites. The anterior is to the right, and posterior is to the left of the video. *kdrl* (vascular endothelial cells, magenta), *fli1* (vascular endothelial cells, nuclear green), Vasa (germ cell, green).

**Movie 3: Z-stacks through the 60 dpf ovary (Figure 3)**. *kdrl* (green), *sox9b* (magenta), DNA (gray). As oocytes mature, they will each be surrounded by at least one vessel.

**Movie 4: Z-stacks through the 60 dpf testis (Figure 4)**. *kdrl* (green), *sox9b* (magenta), DNA (gray). Vessels remain outside of the somatic cells of the testis and surround lobules of maturing sperm.

**Movie 5: 3D rendering of pericytes association with endothelial tip cells (Figure 8)**. Pericytes = *pdgfrb:EGFP* (green), vascular endothelial cells = *kdrl:DsRed* (magenta), DNA = Hoechst (blue)

**Movie 6: 3D rendering and Z-stacks of macrophage engulfing germ cell material (Figure 9C)**. Macrophage = *mpeg:Kaede* (magenta), germ cells *= piwil1:EGFP* (green), DNA = Hoechst (blue).

**Movie 7: Z-stacks of photoconverted macrophage (Figure 9C)**. 24 hours after conversion green and red macrophage could be found in the gonad (blue arrow) while some macrophage in the gonad were only magenta (red arrowhead). In the green channel, we can observe that some macrophage migrate to the gonad within the 24 hour window post-photoconversion (green arrows).

**Movie 8: Movement of a macrophage at 13 dpf over one hour.** Within one hour some macrophage are highly dynamic (arrows) while others remain in place.

## Supplemental Table Legends

**Supplemental Table S1:** Number of individuals counted in each category of cell type being assessed.

**Supplemental Table S2:** Highly expressed genes in each pericyte category: larval pericytes (Cluster 39) (Sheet 1), pdgfrb+ cells (Sheet 2), ovarian non-germ Cells (Sheet 3), ovary pericytes (Sheet 4), and comparisons between each group for the Venn diagram (Sheet 5).

---

**Supplemental Figure S1:**
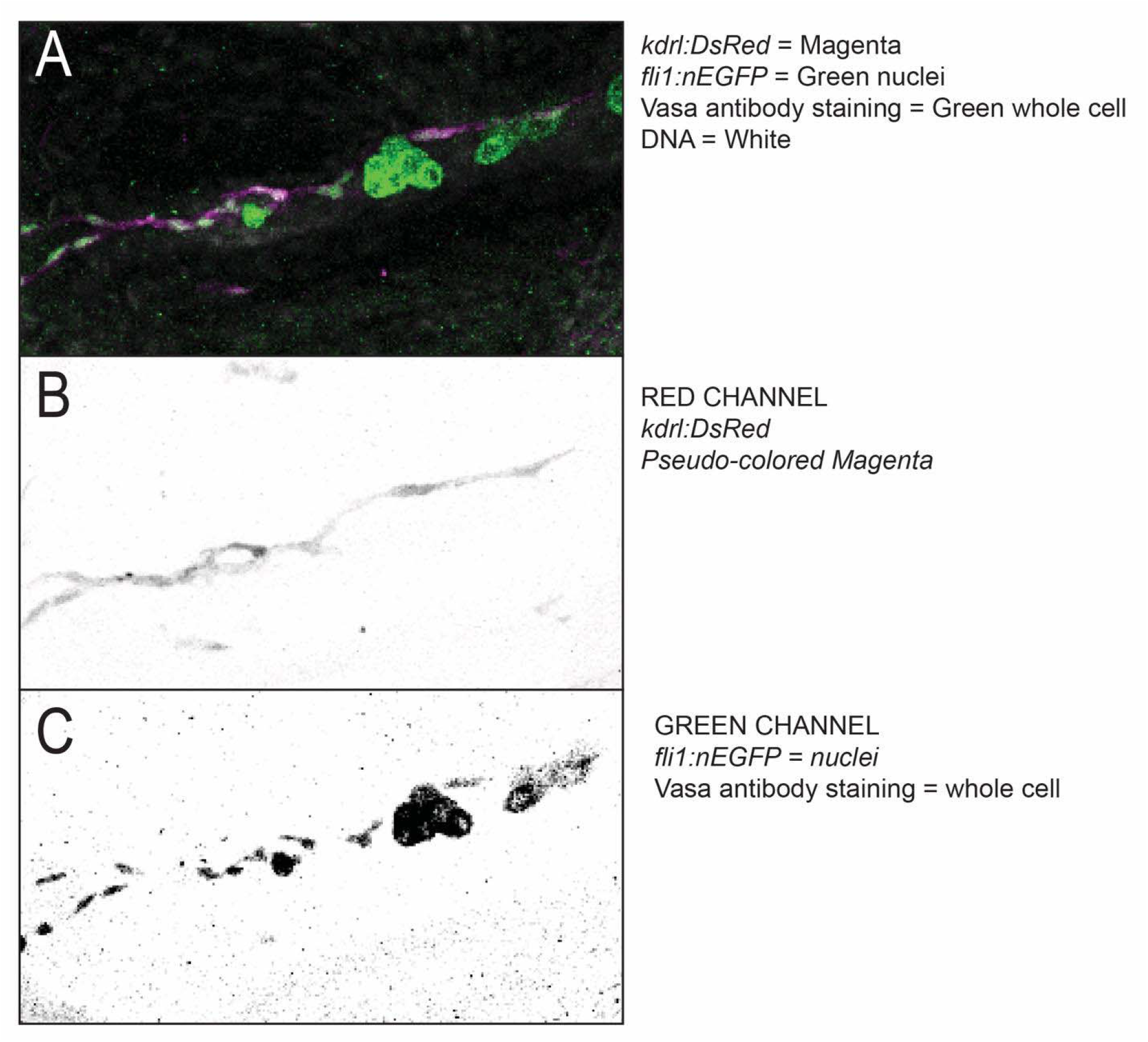
Single channel images of contact between germ cells and vascular endothelial cells.

**Supplemental Figure S2:**
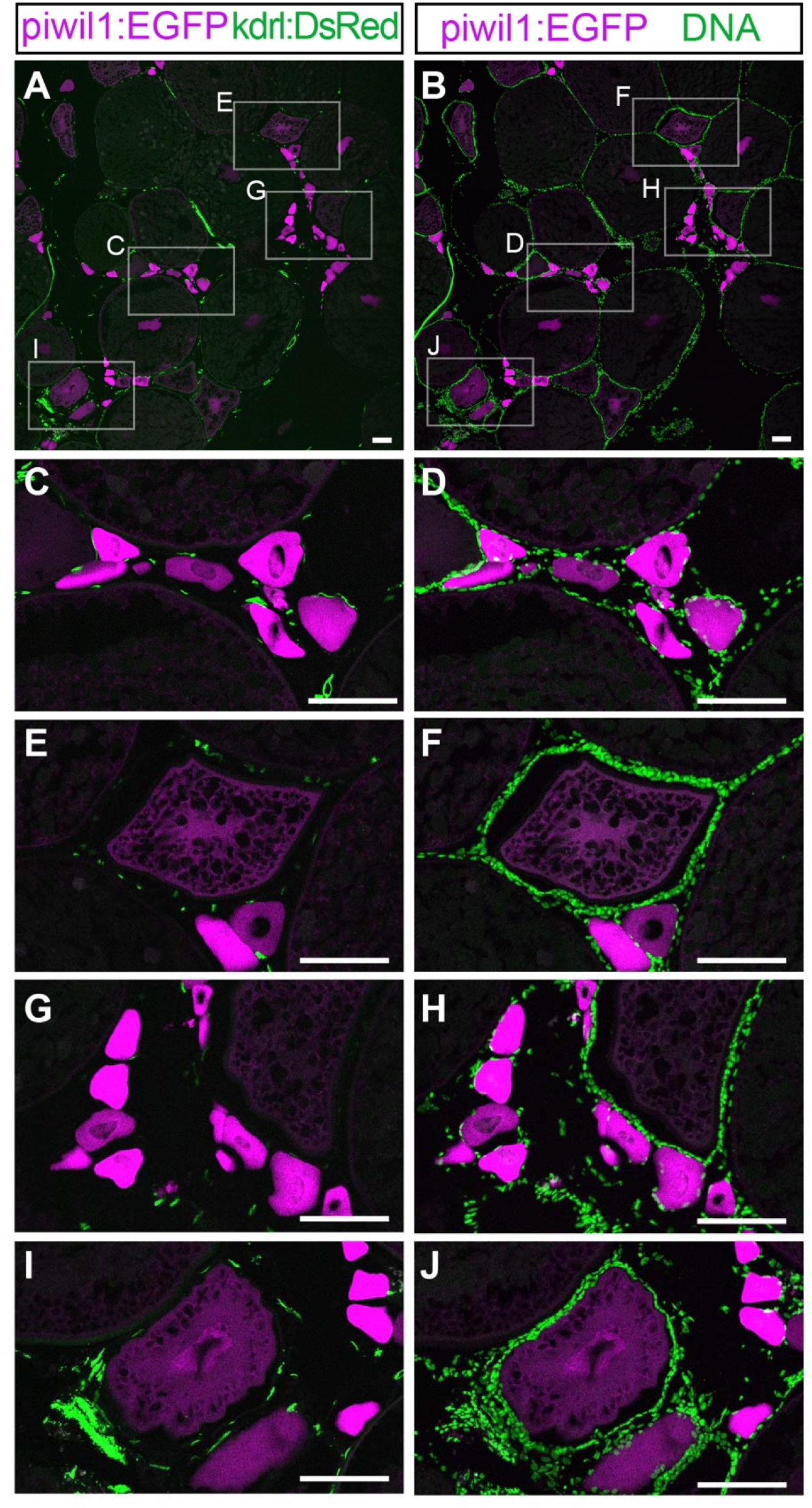
Ovarian vasculature in sectioned tissue. (**A, C, E, G, I**) *kdrl* expressing vessels (green) show the relationship between the vasculature and germ cells (*piwi*, magenta) in the adult zebrafish ovary. **(B, D, F, H, J)**. Hoechst staining (DNA, green) marks the somatic and endothelial cells in the ovary relative to the germ cells (*piwi*, magenta). Note that the more immature oocytes are in indirect contact with the vasculature. Scale: 100 µm.

**Supplemental Figure S3:**
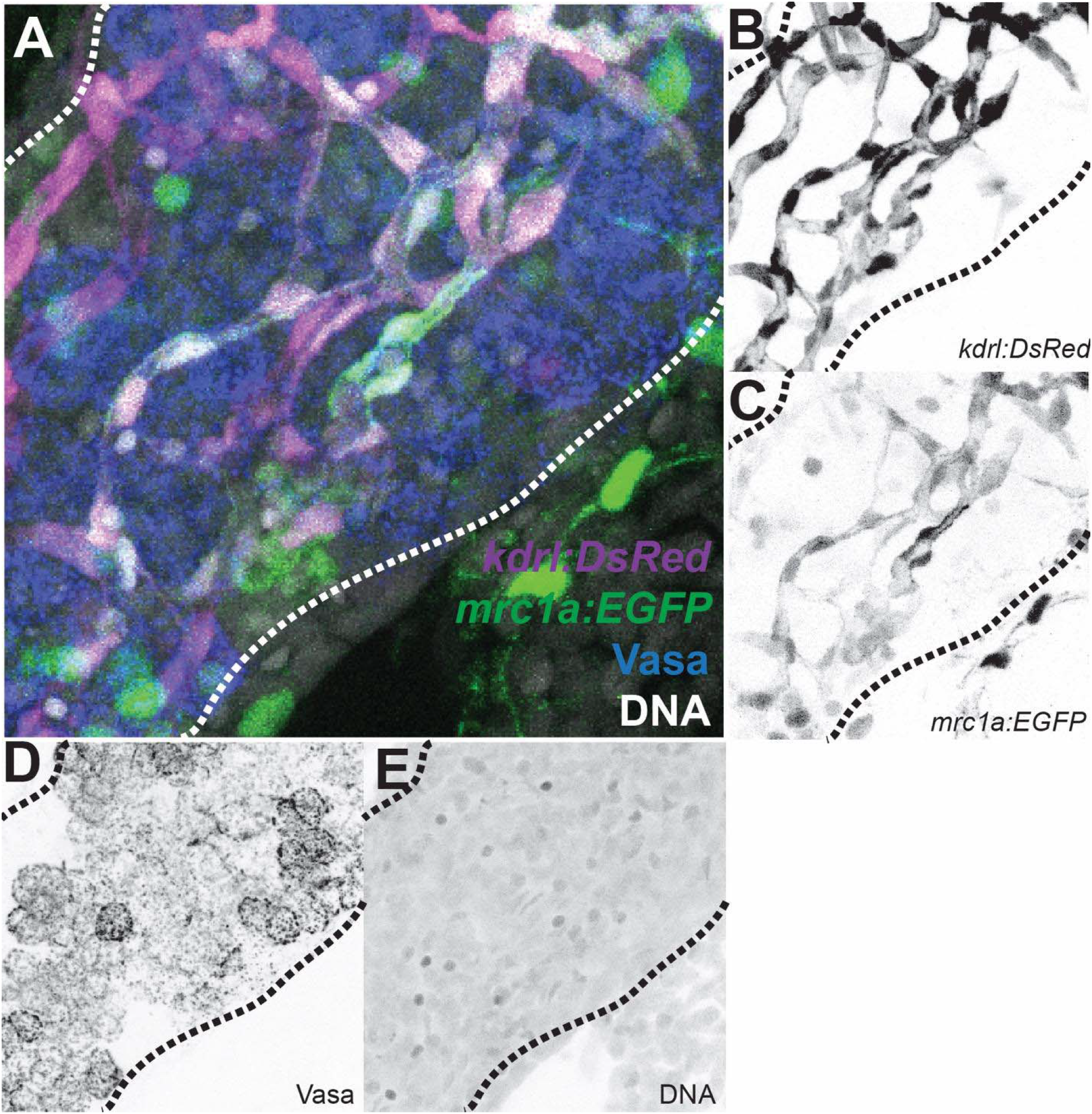
All channels from Figure 8 in of the 20 dpf zebrafish gonad. **A.** vascular endothelial cells= kdrl:DsRed (magenta, panel B), lymphatic endothelial cells = mcr1a:EGFP (green, panel C), germ cells = Vasa (blue, panel D), DNA = Hoechst (white, panel E).

**Supplemental Figure S4:**
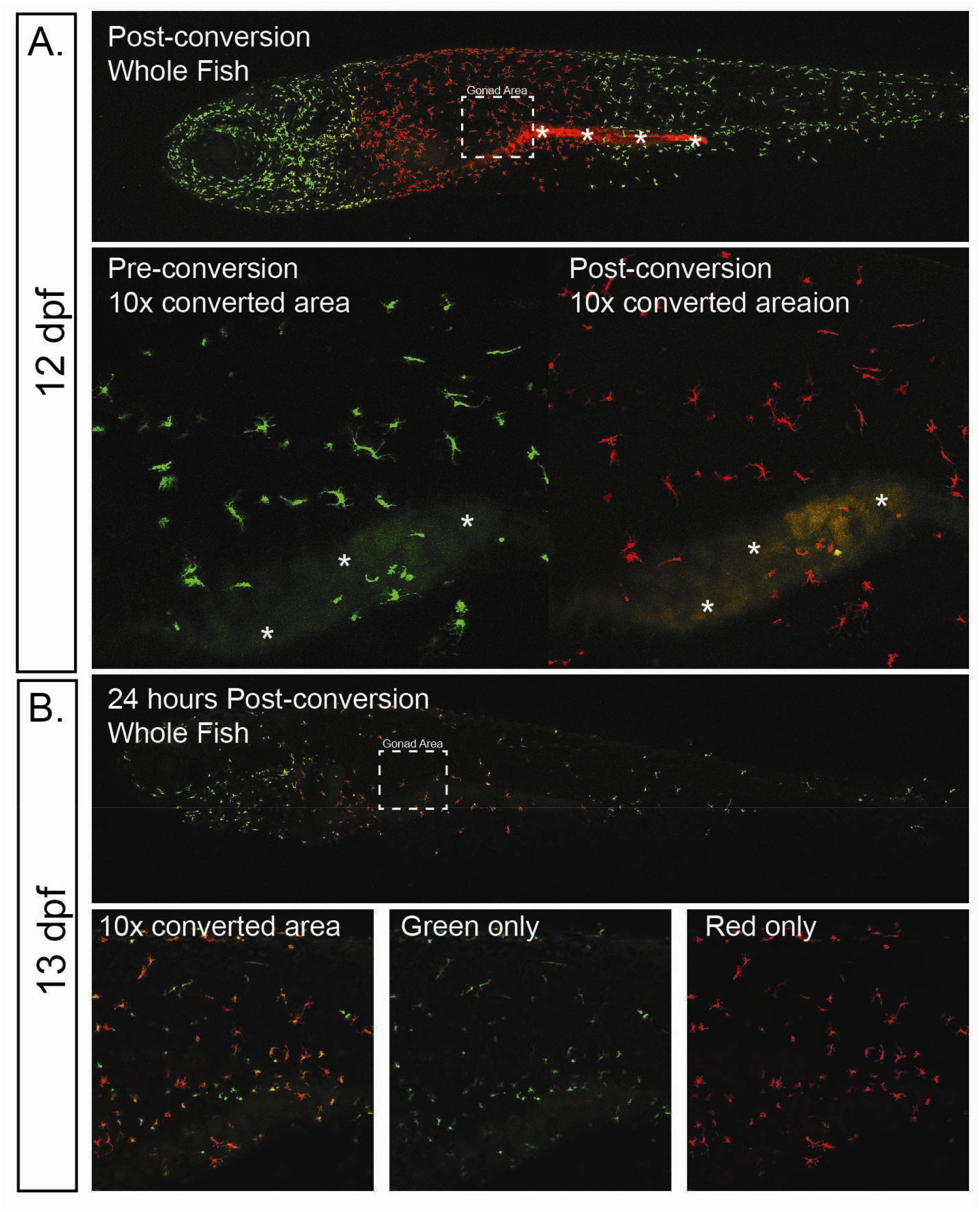
Photoconverted area of *mpeg:Kaede*. **(A)** at 12 dpf the total area which was photo converted and the 10x view of the area pre- and post-conversion. **(B)** After 24 hours, at 13 dpf, the area of *mpeg:Kaede* has expanded and there are both green and red cells present. All images are of different individuals to maximize survival during live imaging. The box denotes the area in which the gonad is located in the body cavity. * denote the gut which is auto-fluorescent both green and red because of the food.

