## Supplemental Table S1 for "Defining the cellular complexity of the zebrafish bipotential gonad": Supplemental Table S1.docx

|  | | **Number of Individuals** | |  |  |
| --- | --- | --- | --- | --- | --- |
| **Stromal Cell Type** | **Dpf** | **Contact** | **No Contact** | **# of independent spawns** | **Standard Length (mm ± st.dev)** |
| VEC | 10 | 0 | 16 (0 Proximal) | 3 | 4.27 ± 0.373 |
|  | 12 | 0 | 13 (4 Proximal) | 4 | 5.08 ± 0.568 |
|  | 15 | 4 | 6 (8 Proximal) | 3 | 6.03 ± 0.539 |
|  | 20 | 15 | 2 | 3 | 7.21 ± 0.666 |
|  | 25 | 15 | 0 | 3 | 8.9 ± 1.21 |
| LEC | 10 | 0 | 9 | 3 | 3.82 ± 0.359 |
|  | 12 | 2 | 15 | 4 | 4.30 ± 0.632 |
|  | 15 | 7 | 6 | 3 | 5.11 ± 0.636 |
|  | 20 | 13 | 0 | 4 | 7.17 ± 1.04 |
|  | 25 | 11 | 1 | 2 | 9.03 ± 1.59 |
| Perivascular  /Pericytes | 10 | 0 | 6 | 2 | 4.429 ± 0.319 |
|  | 12 | 1 | 6 | 2 | 5.271 ± 0.520 |
|  | 15 | 8 | 8 | 3 | 5.72 ± 0.918 |
|  | 20 | 8 | 1 | 3 | 7.61 ± 0.515 |
|  | 25 | 14 | 0 | 3 | 8.5 ± 0.775 |
| Macrophage | 10 | 3 | 9 | 2 | 4.16 ± 0.381 |
|  | 12 | 6 | 8 | 3 | 4.74 ± 0.849 |
|  | 15 | 11 | 2 | 3 | 5.47 ± 0.993 |
|  | 20 | 14 | 0 | 3 | 7.67 ± 1.49 |
|  | 25 | 9 | 0 | 1 | 8.95 ± 0.568 |
